# Development of Potent and Effective Synthetic SARS-CoV-2 Neutralizing Nanobodies

**DOI:** 10.1101/2021.05.06.442911

**Authors:** Maxwell A. Stefan, Yooli K. Light, Jennifer L. Schwedler, Peter R. McIlroy, Colleen M. Courtney, Edwin A. Saada, Christine E. Thatcher, Ashlee M. Phillips, Feliza A. Bourguet, Catherine M. Mageeney, Summer A. McCloy, Nicole M. Collette, Oscar A. Negrete, Joseph S. Schoeniger, Dina R. Weilhammer, Brooke Harmon

**Affiliations:** Systems Biology Department, Sandia National Laboratories, Livermore, CA; Biotechnology and Bioengineering Department, Sandia National Laboratories, Livermore, CA; Biosciences and Biotechnology Division, Lawrence Livermore National Laboratories, Livermore, CA

## Abstract

The respiratory virus responsible for Coronavirus disease 2019 (COVID-19), Severe acute respiratory syndrome coronavirus 2 (SARS-2), has impacted nearly every aspect of life worldwide, claiming the lives of over 2.5 million people globally, at the time of this publication. Neutralizing nanobodies (V_H_H) represent a promising therapeutic intervention strategy to address the current SARS-2 pandemic and provide a powerful toolkit to address future virus outbreaks. Using a synthetic, high-diversity V_H_H bacteriophage library, several potent neutralizing V_H_H antibodies were identified and evaluated for their capacity to tightly bind to the SARS-2 receptor-binding domain (RBD), to prevent binding of SARS-2 spike (S) to the cellular receptor Angiotensin-converting enzyme 2 (ACE2), and to neutralize viral infection. Preliminary preclinical evaluation of multiple nanobody candidates demonstrate that they are prophylactically and therapeutically effective *in vivo* against *wildtype* SARS-2. The identified and characterized nanobodies described herein represent viable candidates for further preclinical evaluation and another tool to add to our therapeutic arsenal to address the COVID-19 pandemic.

**Author Summary:** To fully address the on-going pandemic caused by severe acute respiratory syndrome coronavirus 2 (SARS-2), it will be important to have both vaccines and therapeutic strategies to prevent and mitigate the effects of SARS-2. In this study, we describe the identification and characterization of potently neutralizing humanized single domain heavy chain (V_H_H) antibodies that have binding affinity for both the original Wuhan strain and widely circulating B.1.1.7/UK strain. V_H_H antibodies have the same therapeutic potential as conventional antibodies in half the size and with greater stability and solubility. Using a synthetic humanized high-diversity V_H_H phage library we identified several candidates with strong affinity for the SARS-2 spike that block the interaction of SARS-2 spike with the cellular receptor ACE2, and effectively neutralize infection with SARS-2 *in vitro*. By sequencing viral escape mutants generated in the presence of each V_H_H we mapped the binding sites of the V_H_H antibodies and assessed their affinity against newly emerging SARS-2 variants. Finally, we demonstrate that two of these V_H_H antibodies show prophylactic and therapeutic efficacy *in vivo* against challenge with SARS-2. This study establishes that screening highly diverse V_H_H phage libraries against viral threats can yield highly effective therapeutic agents in real time.

## Introduction

The novel respiratory virus, severe acute respiratory syndrome coronavirus 2 (SARS-2), has infected over 146 million and killed over 3 million people worldwide at the time of this publication(1), causing the worst global health crisis since the 1918 Spanish influenza pandemic. In addition to the cost of lives lost to COVID-19, this virus has wreaked havoc on the global economy and highlighted the threat that emerging diseases pose to global security.(2) It is important not only to work to develop effective vaccines for pre-exposure prophylaxis, but to create new treatments to prevent and mitigate severe disease.

Viral neutralizing antibodies are an effective therapeutic intervention for COVID-19, as the current pandemic response has shown. High titer convalescent plasma has emergency use authorization by the U.S. FDA for the treatment of hospitalized patients early in the disease course and/or with impaired humoral immunity; however, batch-to-batch variability results in various levels of success, limiting its reliability as a treatment.(3) Monoclonal antibody therapies offer improved properties due to their target specificity, uniformity, favorable pharmacokinetics, and their ability to potently block cell entry, the first step of viral infection.(4-7) More recently, promising clinical trial data demonstrating that a single intravenous infusion of monoclonal antibody cocktail significantly reduced COVID-19 related hospitalization and death in comparison to placebo.(8, 9) These results and other clinical trial findings have resulted in emergency use authorization for several cocktails by the FDA for people 12 years and older who test positive for SARS-2 and are at high risk for progressing to severe COVID-19.(8, 9) Methods for improved development and characterization of novel neutralizing antibodies can be part of a critical toolset to combat the COVID-19 pandemic and future epidemics, by providing lower cost, easier manufacturability, and diverse functionality, including response to emerging variants.

Nanobodies (V_H_H) are the variable region of single-domain heavy chain antibodies (sdAbs), which lack the light chain and the CH1 domain, are derived from camelids, and are smaller (∼75 kDa) than human or murine immunoglobulin G (IgG) antibodies (150kDa).(10) V_H_H antibodies are highly soluble, stable, extremely versatile, and have unique structural attributes in their complementarity determining region 3 (CDR3) loop which can facilitate binding to antigen sites inaccessible to traditional IgG antibodies.(11, 12) Neutralization of SARS-2 is achieved by targeting antibodies to the spike (S) receptor-binding domain (RBD), which engages the Angiotensin-converting enzyme (ACE2) receptor to facilitate cell entry. The S protein is heavily glycosylated, limiting viable epitope availability with potential for therapeutic efficacy.(13) Compact V_H_H antibodies may have access to alternate epitopes on the SARS-2 S protein that are sterically inaccessible to traditional antibodies.

Herein, we describe a rapid discovery process for SARS-2 V_H_H antibody therapeutics, going from molecular discovery of high-affinity variable regions to demonstration of therapeutic efficacy of fully humanized V_H_H antibodies in mice infected with SARS-2. We describe the construction of a high-diversity synthetic humanized V_H_H phage library (∼3.2 × 10^10^) that was used to identify several V_H_H antibodies that are high-affinity binders to SARS-2 S protein and RBD. The top V_H_H candidates were produced as human single-domain heavy chain antibodies (hIgG1 Fc) and were evaluated for their ability to block the interaction between purified SARS-2 S and the ACE2 receptor. Effective V_H_H antibodies were then screened for their ability to block infection of Vero cells with SARS-2 pseudotyped vesicular stomatitis virus (VSV-SARS-2-GFP), followed by verification of neutralization of *wildtype* (*WT*) SARS-2 (USA-WA 1/2020). The epitope targets were mapped by sequencing neutralization escape mutants generated in the presence of each V_H_H. The binding affinity of the V_H_H antibodies to RBD with mutations generated in this study or from circulating variants was assessed with kinetic studies using biolayer interferometry (BLI). We also demonstrate that two of these V_H_H antibodies show efficacy *in vivo* both prophylactically and therapeutically against challenge with fully virulent *WT* SARS-2.

## Materials and Methods

### DNA Manipulation and Production of Proteins

pSF-CMV-SARS-2-S was constructed as follows. DNA fragments encoding amino acids 1-1208 of the SARS-2 S gene (GenBank:MN908947.3) were amplified as several independent fragments to introduce several mutations described in Wrapp et. al. from a vector containing the full-length codon optimized S gene.(14) Additionally, a double stranded DNA fragment encoding a C-terminal T4 fibritin trimerization domain, TEV cleavage site, Twin-Step-Tag, and octahistadine was commercially obtained. pSF-SARS-2-RBD was produced by commercial production of double stranded DNA encoding a N-terminal Kozak, a signal peptide (MDWTWRFLFVVAAATGVQS), 319-577 of the SARS-2 S gene (GenBank:MN908947.3), a TEV site, and a C-terminal, decahistadine tag. Mutant SARS-2 RBDs were produced by site-directed mutagenesis. All fragments were synthesized commercially (IDT) and terminal fragments had overlapping sequence with the downstream digested vector. All DNA fragments and pSF-CMV vector restriction digested with *EcoRI* and *BamHI* were assembled using NEBuilder HiFi DNA master.

Constructs for expression of ACE2-Fc (human crystallizable fragment (Fc) domain) fusion proteins were produced as follows. pAce2-huFc was produced by subcloning the ectodomain on human ACE2 (Sino Biological) into pCR-Fc using the *NotI* and *BamHI* restriction sites. To produce Ace2-rbFc (rabbit Fc domain), a gBlock (IDT) of this same region of Ace2 was subcloned into pFUSE rIgG-Fc2.

Soluble SARS-2 S and RBD were both produced by transient expression in Expi293F suspension cells. Cells were transfected with 1 μg/mL plasmid at a density of 3.0 × 10^6^ cells/mL using Expifectamine. Cells were supplemented as instructed by the manufacturer and grown at 37°C with 8% CO_2_. Cell supernatant was harvested on day 4 by centrifugation at 4,000 x g for 30 min at 4°C. Clarified supernatant was passed through a 0.22 μm filter and then applied to a 5 mL HisTrap Excel column preequilibrated with 10 mM Tris (pH 8.0), 300 mM NaCl for SARS-2 S and 20 mM phosphate (pH 7.4), 300 mM NaCl for SARS-2 RBD. The column was washed with 10 column volumes equilibration buffer followed by 5 column volumes equilibration buffer with 20 mM imidazole. The S and RBD proteins were eluted with a step gradient to 500 mM imidazole. Fractions containing SARS-2 S protein were pooled and dialyzed in 20 mM HEPES (pH 8.0), 200 mM NaCl prior to concentration. Protein was then concentrated and filtered prior to application to a Enrich 650 SEC column equilibrated with dialysis buffer for SARS-2 S and PBS for SARS-2 RBD. Purified SARS-2 proteins were stored at -80°C.

ACE2-huFc, ACE2-rbFc, and V_H_H-huFc antibodies were produced by transient expression in CHO-S cells using the ExpiCHO expression system. In brief, cells were transfected at a density of 6 × 10^6^ cells/mL and grown for 18 hours at 37 °C with 8% CO2. Cells were then supplemented as per the manufacture’s guidelines and transferred to 32 °C with 5% CO2. Cell supernatant containing soluble protein was harvested on day 10 by centrifugation at 4,000 x g for 30 min at 4°C. Clarified supernatant was passed through a 0.22 μm filter and the applied to a 1 mL MabSelect PrismA column. The column was washed with 20 mM sodium phosphate (pH 7.4), 150 mM NaCl. ACE2-Fc fusion proteins were eluted with 100 mM sodium citrate (pH 3.0) and immediately neutralized with 1M Tris (pH 9.0). Protein was then concentrated and filtered before application to a Enrich 650 SEC column equilibrated with 10 mM sodium phosphate (pH 7.2), 140 mM NaCl. Purified ACE2-huFc, ACE2-rbFc, and V_H_H-huFc were stored at -80°C. V_H_H-huFc antibodies identified from screening were commercially obtained from GenScript.

### V_H_H Phage Library Construction and Production

Synthetic V_H_H library was designed as follows. A high diversity library was designed by incorporating the natural prevalence of amino acids at positions in CDR1 and CDR2 based off 670 functional V_H_H antibodies deposited on sdAb-DB (www.sdab-db.ca).(15) Amino acids cysteine and methionine were omitted from all CDR loops and full diversity was used for CDR3. Asparagine was also omitted from CDR1 and CDR2. The library was also designed such that there would be 3 different lengths of CDR3 (9-, 12-, and 15-amino acids). Linear double-stranded DNA fragments incorporating the specifications mentioned above as well as terminal flanking BglI restriction sites were synthetically produced commercially by Twist Bioscience. A humanized V_H_H framework characterized in Moutel et. al. was used to house the designer CDRs.(16)

Library was assembled as follows. The linear library DNA fragment was amplified by PCR. PCR reaction was divided into a 384 well format to minimize potential amplification bias. Five cycles of PCR were performed (Step 1: 98°C, 3 minutes, 1 cycle; Step 2: 98°C, 10 seconds; 68°C, 10 seconds; 72°C, 15 seconds, 5 cycles; Step 3: 72°C, 10 minutes, 1 cycle; Step 4: 12°C, hold). Amplified DNA was pooled and purified by Monarch Nucleic Acid Purification Kit. The library was digested with 5U/μg DNA SfiI at 50°C for 16 hours. Digest reactions were again column purified as previously mentioned. Ligations were set up as follows. pADL20c (205 μg) previously digested with BglI and treated with rSAP was added to a 40 mL reaction containing the linear digested library (42 μg) at a vector to insert ratio of 1:2, 1 mM ATP and 1,100,000 units of T4 Ligase. The reaction was allowed to proceed at 16°C for 16 hours, then at 37°C for 1 hour, and lastly was heat inactivated at 70°C for 20 minutes. DNA was purified and concentrated by a modified ethanol precipitation protocol utilizing tRNA carrier at 15 μg/mL.

Transformation of the library into TG1 *E. coli* was performed as follows. A total of 9 liters of fresh electrocompetent TG1 *E. coli* cells were produced and 150 electroporations performed. To 350 μL cells, 15 μL of ligated DNA was added and electroporation performed. After 1-hour recovery with the addition of 650 μL SOC at 37°C, cells were plated on 2xYT agar plates supplemented with 100 mM glucose and 100 μg/mL carbenicillin (2xYT-GA). *E. coli* was harvested after growth at 37°C for 16 hours, supplemented with glycerol to 10%, and stored at -80°C in aliquots for further use. Diversity of the nanobody library was determined by both NGS and colony PCR.

The V_H_H library was added to 2xYT-GA to a final OD_600_ of 0.08 and allowed to grow at 37°C until an OD_600_ of 0.5 was reached. Superinfection with CM13 (ADL) was performed with 2.0 × 10^12^ helper phage per liter of *E. coli* culture for 15 minutes without shaking followed by rigorous shaking for 30 minutes. Cell were collected by centrifugation at 5,500 x g for 10 minutes and resuspended in the previous volume used of 2xYT supplemented with 100 μg/mL carbenicillin and 50 μg/mL kanamycin (2xYT-AK). *E. coli* were grown overnight with shaking at 28°C.

Supernatant containing packaged phagemid was clarified by centrifugation at 6,000 x *g* for 15 minutes and supplemented with one-fourth volume 20% PEG-8000 and 2.5 M NaCl. Phage were precipitated overnight at 4°C and were collected by centrifugation at 5,500 x g for 60 minutes at 4°C. The pellet containing phage was resuspended in PBS and centrifugated for 15 minutes at 5,500 x g to remove *E. coli* particulates. Clarified phage was again precipitated as mentioned above and incubated on ice for 60 minutes. The phage was finally centrifuged at 17,900 x g for 10 minutes at 4°C, the supernatant was removed, and the pellet was again centrifuged at 17,900 x g for 1 minute to remove trace amounts of PEG-8000. The pellet was resuspended in PBS and passed through a 0.22 μm filter before determining the colony forming units (CFU).

### Next-Generation Sequencing

All samples for NGS were processed as follows. The minimum region containing all 3 CDR domains, approximately 300 bps, was excised from the pADL20c backbone by two-step restriction digests, *BglI* followed by *DdeI*/*BstEII* double digests on the gel-purified small fragment from the *BglI* restriction reaction. The sequencing library was prepared using NEBNext^®^ Ultra™ II DNA Library Prep Kit for Illumina^®^(NEB) and xGen™ UDI-UMI Adapters (IDT) as instructed in the manufacture’s manuals and sequenced on Illumina NextSeq 500/550 platform with High Output v2.5 300-cycles, paired-end mode. BCL files were converted to FASTQ and demutiplexed using the bcl2fastq script from MyIllumina (https://my.illumina.com). The quality filtering and adaptor trimming were performed using fastp (https://github.com/OpenGene/fastp) with following parameters, -q 30 -l 100 -x 7. R2read was reformatted to be reverse complemented and merged with R1 read using BBTools (BBMap, https://sourceforge.net/projects/bbmap/).(17) Three CDR domains with correct sequence lengths were extracted, concatenated and translated and counted unique amino acid sequences using a custom python script.

### Affinity Selection for SARS-2 S Clones by panning

A total of four rounds of affinity selection were conducted, three rounds against soluble full-length SARS-2 S and a final round against SARS-2 RBD. Panning was performed by immobilizing antigen in 96-well Immulon® HBX microplates. For the first round of panning 3 μg SARS-2 S was immobilized in 50 μL coating buffer (100mM NaHCO_3_ (pH 8.3), 150 mM NaCl) overnight at 4°C. Wells containing antigen were washed five times with 300 μL PBS-t (phosphate buffered saline supplemented with 0.05% Tween-20). Wells were blocked with 300 μL Pierce PBS Protein Free Blocking Solution for 2 hours. After another round of washing, 200 μL of 2.5 × 10^12^ CFU/mL phage were added to each well in blocking buffer and allowed to incubate with vigorous shaking at 25°C for 2 hours. Wells were extensively washed with increasing amounts of Tween-20, with a peak concentration of 0.5% Tween-20. Phage were eluted with the addition of 50 μL of elution buffer (10 mM Tris pH 7.4, 137 mM NaCl, 1 mM CaCl_2_ and 100 µg/mL Trypsin) by incubating for 30 minutes at 37°C. Eluate was removed and pooled and a second 50 μL of elution buffer was added for 20 minutes. This was collected and pooled with the first round of elution buffer. Eluted phage was added to an equal volume of log phage TG1 *E. coli* for reamplification and quantification.

For all subsequent rounds of panning, 2 μg of antigen was used to coat wells. For round 2, a total of 10^12^ CFU phage were used for panning. For rounds 3 and 4, panning was conducted with an input of 10^11^ CFU phage. With each round of panning, washing was progressively increased from 12 washes in the first round to 15, then to 21 for the final two rounds. For all rounds after the first, a maximum of 1% Tween-20 used for washing. The plates were then extensively washed with PBS to remove excess Tween-20. Additionally, the blocking solution for each round of panning was cycled. For the second round PBS-t supplemented with 1% bovine serum albumin (BSA) (w/v) was used, for the third round PBS-t supplemented with 2% non-fat milk, and for the final round BSA was again used. A denaturing step was conducted before the last round of panning against the SARS-2 RBD to remove unstable clones by heating the phage to 70°C for 15 minutes before panning. All other elution, reamplification, and tittering steps were conducted as mentioned above.

### Polyclonal and Monoclonal Phage ELISA

Polyclonal phage ELISA was used to characterize antigenic enrichment and specificity for each round of panning. Multiple antigens were tested including BSA, SARS-CoV-1 S, SARS-2 S, and SARS-2 RBD. Immulon HBX microtiter 384 well plates were coated with at 0.05 μg/well of each antigen overnight at 4°C. Plates were washed 5 times with PBS-t and blocked with 50 μL of PBS with 2% non-fat milk and 0.2% Tween-20 (MPBS-t). Plates were washed and 10^9^ phage from the unenriched library and each round of panning was added to each well for all of the antigens in blocking solution. After 2 hours of shaking at 25°C the plates were again washed five-times and mouse anti-M13 coat protein (Thermo) was added 1:2000 in MPBS-t for 1 hour. The plates were again washed, and rabbit anti-mouse IgG HRP (Thermo) was added 1:1000 in MPBS-t for 1 hour with shaking. The plates were finally washed 5 times with PBS-t and developed with TMB Ultra substrate (Thermo) and quenched with 2 M H_2_SO_4_ before reading the absorbance at 450 nm. Monoclonal phage ELISAs were performed in a similar fashion to the polyclonal ELISA mentioned above with homogeneous preparations of V_H_H-phage to validate individual clones.

### V_H_H-huFc ELISA

Indirect ELISAs were performed as follows. Full-length soluble SARS-2 S or RBD were immobilized on Immulon 384-well microtiter plates (0.05 μg/well) overnight at 4°C in coating buffer. After washing, plates were blocked for 2 hours with Pierce PBS Protein Free Blocking buffer (Thermo). Serial dilutions of V_H_H-huFc antibodies in blocking buffer were added to plates for 2 hours after washing. Secondary antibody, goat anti-human H+L IgG HRP (Thermo) was added for an additional hour before washing. Plates were developed with TMB Ultra (Thermo) and the reaction stopped by addition of equal volume 2 M H_2_SO_4_ after washing. Absorbance was read at 450 nm.

Competition of soluble ACE2-huFc by V_H_H-huFc candidates identified from display screening was performed as follows. SARS-2 antigen was immobilized and blocked as described above. After blocking, serial dilutions of V_H_H-huFc antibodies were prepared in blocking solution and allowed to incubate for at least 2 hours, after which ACE2-rbFc was added to a final concentration of 0.1 μg/mL (SARS-CoV-1 S) or 0.06 μg/mL (SARS-2 S) and allowed to incubate for 1 hour with shaking at 25°C. Plates were washed and 1:10,000 HRP-conjugated goat anti-rabbit IgG (Thermo) was added. Plates were developed as described above.

### Differential Scanning Fluorimetry

Thermostability for the V_H_H-huFc antibodies was determined by differential scanning fluorimetry (DSF). Each reaction contained 10 μg of V_H_H-huFc in PBS with 10 μM SYPRO Orange (ThermoFisher). Reactions were heated from 10-95°C at a heating rate of 1°C/min and monitored in the FRET channel of a BioRad CFX96. The melting point for each of the V_H_H-huFc antibodies was calculated by the first derivative method.(18)

### Viruses and Cell Culture

Vero and Vero E6 cells (African green monkey kidney, ATCC CCL-81 and CRL-1586, respectively) were maintained in culture medium supplemented with 10% fetal bovine serum (FBS), 100 units/ml penicillin, and 100 μg/ml streptomycin (Invitrogen), at 37 °C in 5% CO2. For risk group (RG) 2 experiments and viral stocks, Vero cells were cultured in supplemented minimum essential medium alpha (alpha MEM). For RG 3 experiments and viral stocks, Vero E6 cells were cultured in supplemented Dulbecco’s Modified Eagle’s Medium (DMEM). A pseudotyped, replication-competent Vesicular stomatitis virus (VSV) expressing eGFP as a marker of infection and the SARS-2 spike gene (VSV-SARS-2) in place of its own VSV-G gene was provided by Dr. Sean Whelan(19, 20). Pseudotyped, single cycle VSV particles displaying the Severe acute respiratory syndrome (SARS-1) spike protein (VSV-SARS-1-GFP) were derived from recombinant VSV-DG-GFP in which the VSV-G envelope protein has been replaced with GFP. VSV-SARS-1-GFP single cycle particles were produced in Expi293 cells through transfection per manufacturer’s instructions with the SARS-CoV-1 spike expression plasmid (Sino Biological, Cat #VG40150-G-N) containing a 19-aa deletion in the cytoplasmic tail. The Expi293 cells were subsequently infected with VSV-DG-GFP, itself pseudotyped with VSV-G, at 72 h post-transfection using a multiplicity of infection (MOI) of 3. The resulting VSV-SARS-1-GFP pseudotyped viruses were collected at 24 h post-infection. *WT* SARS-2 was deposited by the Centers for Disease Control and Prevention and obtained through BEI Resources, NIAID, NIH: SARS-Related Coronavirus 2, Isolate USA-WA1/2020, NR-52281. An infectious clone of SARS-2 expressing a NeonGreen reporter gene (icSARS-CoV-2-mNG, or SARS-2-NG) was provided by Dr. Pei Yong Shi.(21) Viral stocks were amplified in Vero E6 cells, supernatants were harvested upon extensive cytopathic effect, 48 (VSV-SARS-2-GFP) or 72 (SARS-2) hours post infection. Cellular debris was removed by centrifugation, and aliquots were stored at -80°C. Titers of viral stocks were determined by plaque assay. All work with *WT* SARS-2 was performed in Institutional Biosafety Committee approved BSL-3 and ABSL-3 facilities at Lawrence Livermore National Laboratory using appropriate personal protective equipment (PPE) and protective measures.

### SARS-2 Fluorescent Reporter Neutralization Assays

Serial dilutions of V_H_H-huFc were prepared in supplemented media at 2x the desired final concentration. SARS-2-NG, VSV-SARS-2-GFP, or VSV-SARS-1-GFP was added to the serially diluted V_H_H-huFc antibodies for a final multiplicity of infection (MOI) of 0.2 (RG 2) or 0.1 (RG 3) and allowed to incubate at 37°C for 30 minutes to 1 hour with shaking prior to transfer of virus-V_H_H-huFc mixture to cells seeded in 96 (RG 3) or 384 well plates (RG 2). V_H_H-huFc-virus complexes were incubated with Vero cells at 37°C with 5% CO_2_ for 12-16 h. For RG 2 experiments, cells were subsequently fixed in 4% paraformaldehyde (Millipore Sigma) containing 10 mg/mL Hoechst 33342 nuclear stain (Invitrogen) for 30 min at room temperature, when fixative was replaced with PBS, and images were acquired with the Tecan Spark^®^ Cyto multi-mode plate reader and image cytometer in both the DAPI and FITC channels to visualize nuclei and infected cells (i.e., eGFP-positive cells), respectively (4X objective, covering the entire well). Images were analyzed using the Spark Control Image Analyzer Software. For RG 3 experiments with, cells were lysed in RIPA buffer plus Halt protease inhibitor cocktail (ThermoFisher) and fluorescence was measured using 485 nm excitation and 528 emission wavelengths. Fluorescent values were background subtracted using no-infection controls and normalized to no-treatment infection values. The dose response curves and 50% effective inhibitory concentrations (EC_50_) were generated using Graphpad Prism 9.

### Plaque Neutralization Assay

V_H_H-huFc-*WT* SARS-2 virus complexes were preincubated at 37°C with 5% CO_2_ for 1 hour prior to addition to subconfluent Vero E6 cells in 12 well plates. After a 30-minute incubation of V_H_H-huFc-virus complexes with Vero E6 cells at 37°C with 5% CO_2_, overlays of 2 mL per well of 0.6% microcrystalline cellulose (MCC, Sigma 435244), in 8% FBS, 1% P/S complemented 2X MEM were added. To stain, the MCC was aspirated, wells rinsed with PBS, and 0.4% crystal violet in 100% methanol was added for 10 minutes and then removed. Wells were then washed twice with water, and titers were recorded in plaque forming units (PFU)/mL.

### Biolayer Interferometry

Affinity measurements for V_H_H-huFc antibodies were performed using biolayer interferometry (BLI) using an Octet 384 Red system (Sartorius). Measurements were conducted in 10 mM phosphate (pH 7.4), 300 mM NaCl, 1 mg/mL BSA, 0.1% NP-40. V_H_H-huFc ligands were immobilized on human Fc capturing sensors. *WT* and mutant SARS-2 RBD were used as the analyte and sensograms were fit to a global 1:1 fit. Sensograms which deviated from a 1:1 global fit or fit well to a heterogeneous ligand model were designated as biphasic.

### Generation and Analysis of Escape Mutants

Mutant VSV-SARS-2-GFP virus with the ability to escape V_H_H-huFc neutralization was generated in a similar fashion to that described in Baum et. al.(22) In brief, a series of 500 μL dilutions of V_H_H-huFc antibodies were incubated with 500 μL of 1.5 × 10^6^ PFU VSV-SARS-2-GFP virus for 1 hour at 37°C. The V_H_H-huFc and virus mixture was added to Vero cells and allowed to incubate for up to 72 hours until apparent cytopathic effect (CPE) or widespread viral replication could be observed by GFP fluorescence. Supernatant from wells containing the highest concentrations of V_H_H-huFc was collected and viral RNA isolated from total RNA using TRIzol. A portion of this supernatant was diluted 500-fold and exposed to a series of higher concentrations of V_H_H-huFc antibodies for 1 hour at 37°C prior to addition to Vero cells. Cells were monitored for 60 hours for CPE and GFP fluorescence. Supernatants at the highest concentration of V_H_H-huFc with clear viral replication were collected and processed as mentioned above and viral RNA analyzed as described below.

VSV-SARS-2 genomic RNA was isolated for next-generation sequencing (NGS). For library preparation, 25-100 ng of total RNA was used to deplete rRNA using DNA probes and RNase H provided in RiboErase (HMR) kit (Roche). The depleted RNA was fragmented prior to cDNA synthesis followed by Illumina adaptor ligation using KAPA RNA HyperPrep kit (Roche). The library was analyzed using High Sensitivity DNA ScreenTape 1000 (Agilent) for quality and quantification and pulled together for the final library denaturation. Sequencing was performed on Illumina NextSeq 500/550 platform with High Output v2.5 75-cycle mode. BCL files were converted to FASTQ and demutiplexed using the bcl2fastq script from MyIllumina (https://my.illumina.com). The quality filtering and adaptor trimming were performed using fastp with -q 30 option. The filtered reads were mapped to Spike protein coding region of the SARS-2 Wuhan Hu-1 isolate (GenBank:MN908947.3) using Bowtie 2.(23) The variant calling on the mapped reads was performed using mpileup from Bcftools (https://samtools.github.io/bcftools/) and inspected using Integrative Genomics Viewer (IGV).(24)

### Animal Studies

All animal work was performed in accordance with protocols approved by the Lawrence Livermore National Laboratory Institutional Animal Care and Use Committee. Groups of male and female K18-hACE2 C57BL/6J transgenic mice (Jackson Laboratory) ranging in age from 12-16 weeks were inoculated intranasally with 2.5×10^4^ PFU SARS-2 (USA-WA 01/2020) while under anesthesia (4-5% isoflurane in 100% oxygen). Animals were dosed prophylactically (−24 hours pre-infection) or therapeutically (+24 hours post-infection) by intraperitoneal injection with 10 mg/kg of V_H_H-huFc or isotype control antibody. Body weight was measured daily and any animal falling below 80% of their starting weight was humanely euthanized in accordance with animal welfare guidelines.

### Statistical analysis

Raw data for infection assays measured by reporter fluorescence were compared using a two-tailed t test for each individual experiment. For plate assays, untreated, infected controls and uninfected controls were included on every plate. Antibody-treated cells that were infected were compared to control samples infected with the same virus. Kaplan-Meier survival curves were generated based on two independent experiments and log rank tests were performed with Bonferroni multiple comparison correction applied (GraphPad Prism). P values were considered significant when they were <0.05 (*) and very significant when they were <0.01 (**), <0.001 (***), or <0.0001 (****).

## Results

### Library Construction

The V_H_H library incorporated both the diversity and prevalence of amino acids at key positions in each of the CDR1 and CDR2 derived from a database of 670 functional V_H_H antibodies (**Fig. 1A**).(15) For CDR3, full amino acid diversity and three different lengths were used, 9-, 12-, and 15-amino acids. The length for each CDR 1 and 2 was based on the predominant lengths of these loops observed in the functional V_H_H database. (**S1 Fig)**. Cysteines and methionines were omitted from all CDRs. The library was synthetically produced and the observed prevalence of amino acids at each of the CDR loops paralleled the desired prevalence as determined by NGS (data not shown).

**Fig. 1:**
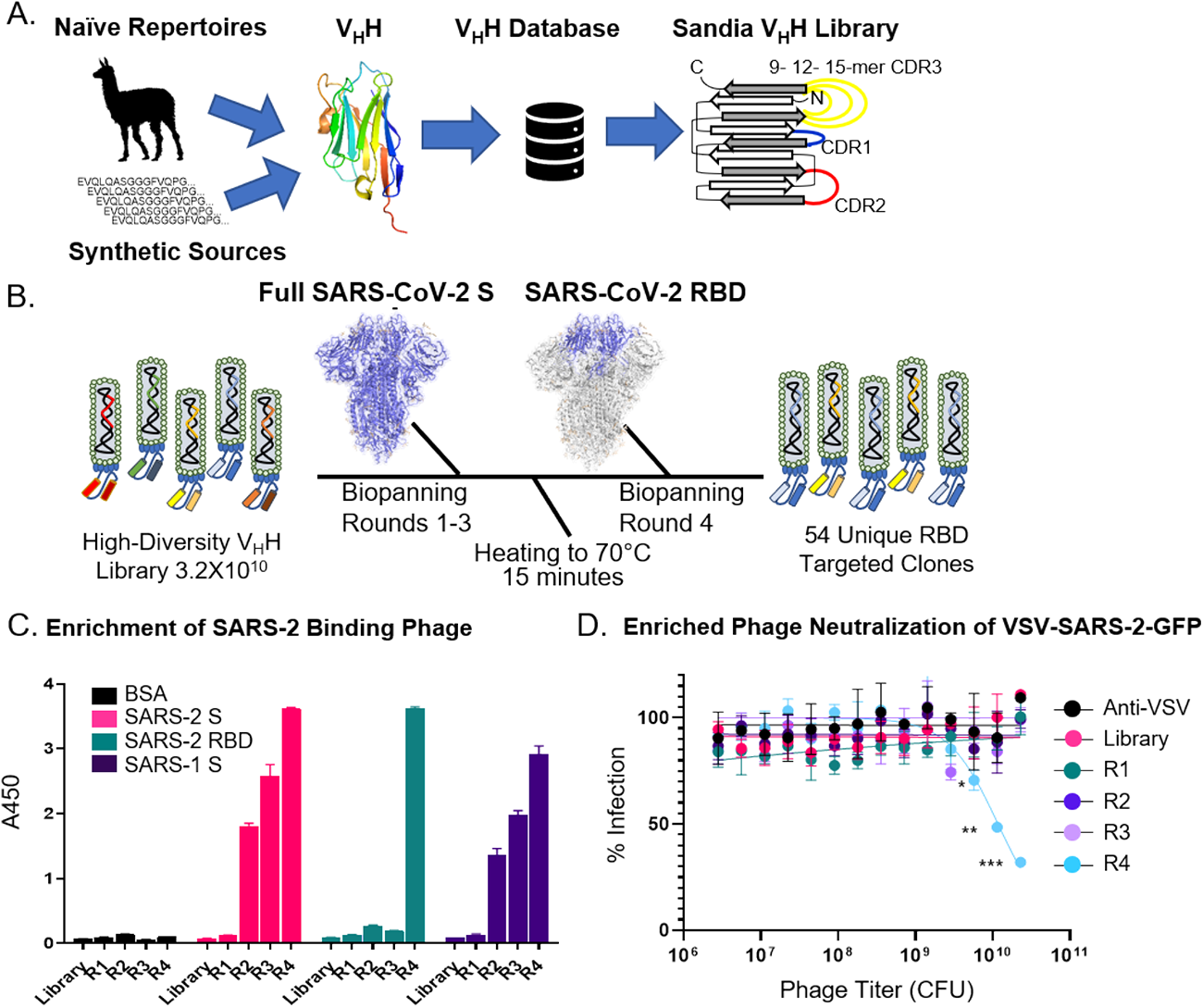
Development of high-diversity V_H_H library and screening campaign for SARS-2 neutralizing V_H_H’s. **A)** Library design using deposited sequences of functional V_H_H’s from various sources. Data was mined to determine optimal CDR length and amino acid prevalence at each amino acid position within each respective CDR. **B)** Schematic for the biopanning approach used to identify neutralizing V_H_H’s directed to the SARS-2 RBD. **C)** Polyclonal ELISA against multiple antigens show enrichment of a positive SARS-2 binding population of displayed V_H_H with each sequential round of panning. Data is from experimental conditions performed in triplicate; the error is the standard deviation from the mean. **D)** Neutralization of VSV-SARS-2 GFP viral infection with polyclonal V_H_H displaying phage from each round of panning. Data is from experimental conditions performed in triplicate; the error is the standard deviation from the mean (***, P < 0.001; **, P < 0.01; *, P < 0.05). The data represent one of 3 experiments with similar results.

In order to obtain sufficient diversity coverage for the library (i.e. transformants), 150 electroporations were performed yielding approximately 3.38 × 10^10^ transformants. To determine the level of success for the ligation of the library into the vector backbone, colony PCR was performed. Of the 408 colonies selected, 395 contained the correct size amplified DNA fragment (95.9%). This value was used to adjust the calculated value for library diversity to 3.24 × 10^10^. Finally, library diversity, quality, and the distribution of CDR3 lengths were assessed by NGS from a total of 39,870,360 reads (**S1 and S2 Table**). The 9-amino acid CDR3 was the most prevalent at 40%, followed by 12-amino acid CDR3 at 34%, and lastly the 15-amino acid CDR3 at 25% of the observed diversity.

Overall, there was good coverage of all represented CDR3s. Approximately 1% of sequences contained a stop codon and 99% of reads were unique sequences (38,592,027 reads). Roughly 1% of reads were duplicates, and 0.01% (1,095 sequences) were present in triplicate. With these corrections the adjusted diversity for this nanobody library is 3.18 × 10^10^.

### Panning against SARS-2 S and RBD

Four rounds of biopanning were used to identify clones that bind to the SARS-2 S protein RBD. For the first three rounds, full-length soluble purified SARS-2 S protein was used, ensuring that conformational integrity of the RBD was maintained for initial selection. A 15-minute heat denaturing step at 70°C was used to remove unstable sequences and a final round of biopanning against SARS-2 RBD was conducted to identify therapeutically relevant V_H_H antibodies (**Fig. 1B**). Enrichment of phage against SARS-2 S was observed over the initial 3 rounds of biopanning (**Fig. 1C** and **S3 Table**) and there was a significant loss in phage recovered when the antigen was shifted to RBD (0.0005% compared to 0.004%).

Polyclonal ELISA showed that over each round of biopanning there was enrichment for SARS-2 S binders. Interestingly, there was relatively little SARS-2 RBD binding until panning was conducted against RBD specifically, indicating preferred epitopes are outside the RBD (**Fig. 1C**). Additionally, neutralization of SARS-2 pseudotyped vesicular stomatitis virus encoding eGFP (VSV-SARS2-GFP)(19) was only observed after RBD biopanning (R4) showing enrichment for RBD binders is required to remove clones binding to epitopes outside the RBD and to identify virus neutralizing clones (**Fig. 1D**). Monoclonal characterization was performed using ELISA, 222 of the 384 clones were designated as hits. Clones that had OD_450_ ≥ 2.5 in the phage coat monoclonal ELISA against both SARS-2 S and RBD and that did not bind BSA were designated as hits (**S2 Fig**.). Of those designated as hits, 54 sequences were identified as unique.

### Evaluation of V_H_H-huFc Candidates

Humanized V_H_H antibodies were produced as a fusion to the hinge region and crystallizable fragment (Fc) domain of a human IgG1, which combines the advantages of the V_H_H with the improved half-life and effector functions of human IgG while reducing the overall size by half that of a conventional antibody(12). All 54 unique V_H_H sequences were commercially produced in a V_H_H-huFc (huIgG1 Fc) format by GenScript for further characterization and triage (**Fig. 2A**). Of the 54 V_H_H-huFc antibodies, 49 showed positive binding to both SARS-2 S and RBD with little to no binding to BSA by ELISA (**S3 Fig**). Those which showed positive binding by ELISA were then tested for their ability to compete with ACE2-huFc for binding to SARS-2 S by a competition ELISA (**S4 Table**). Finally, 46 candidates were evaluated for their ability to neutralize VSV-SARS-2-GFP infection *in vitro*, with 34 V_H_H-huFc antibodies demonstrating full or partial neutralization of infection (**S4 Table**).

**Fig. 2:**
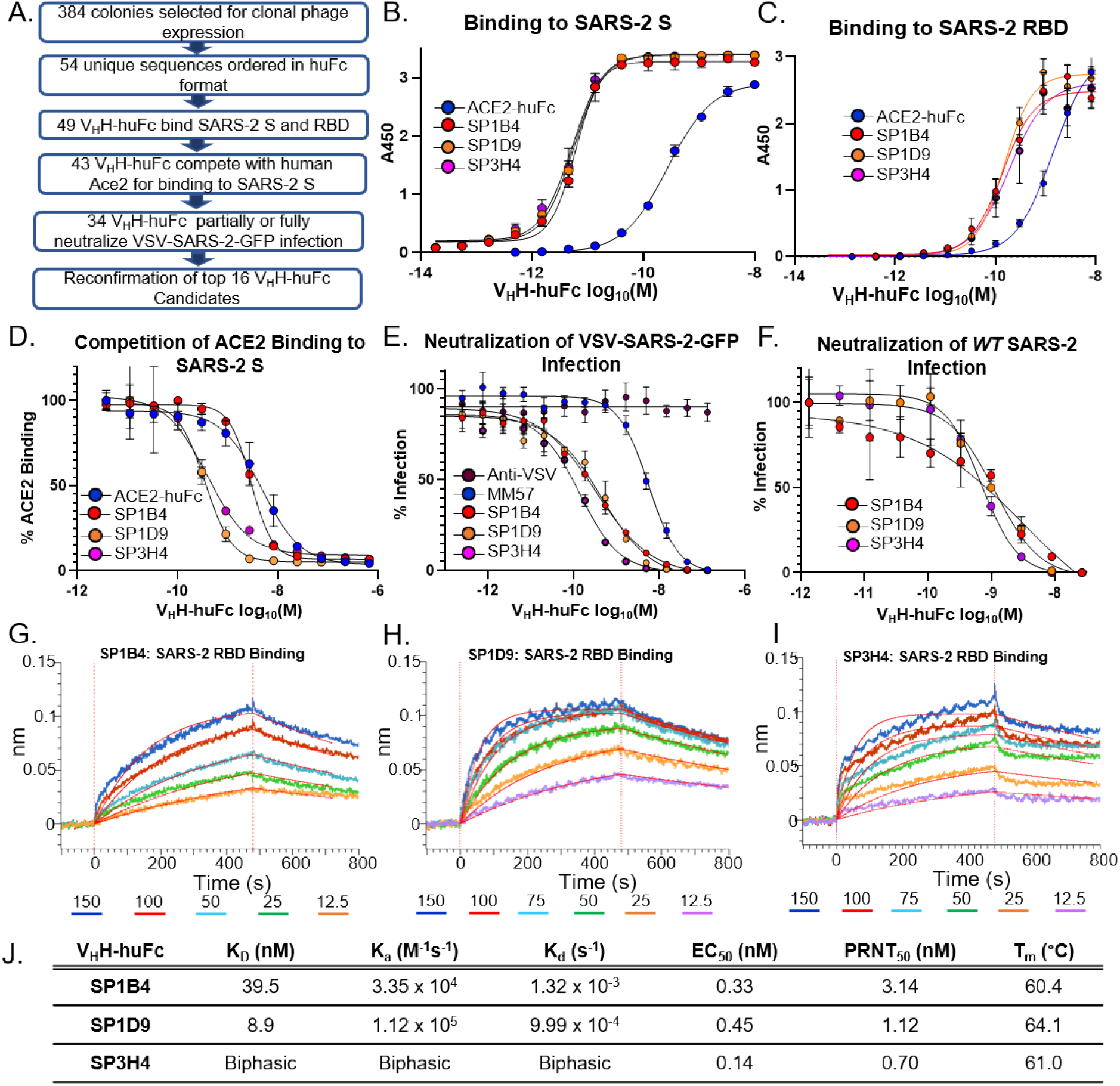
Evaluation of V_H_H-huFc candidates. **A)** Triage flowchart for selection of top candidates, from clonal phage ELISA to selection of 16 candidates for reconfirmation. **B)** Reconfirmation ELISA of top three candidates binding to SARS-2 S and **C)** SARS-2 RBD. **D)** Top candidate V_H_H-huFc’s compete with ACE2-huFc for binding to SARS-2 S by competition ELISA and **E)** neutralize VSV-SARS-2 GFP infection of Vero cells. The data for a-e are from experimental conditions performed in triplicate, the error is the standard deviation from the mean. **F)** All three V_H_H-huFc’s also neutralize *WT* SARS-2 in a plaque reduction neutralization assay. The data from this is the mean of the plaque assay performed in duplicate, the error is the standard deviation from the mean. BLI sensograms show binding to SARS-2 RBD by SP1B4 **(G)**, SP1D9 **(H)**, and SP3H4 **(I)**. At least 5 concentrations were used for global fit analysis. **J)** Table summarizing kinetic properties, neutralizing efficacy, and thermal stability properties for SP1B4, SP1D9, and SP3H4. Melting temperatures from DSF were determined from two independent experiments each performed in triplicate.

For reconfirmation of binding and neutralization, 16 V_H_H-huFc antibodies were selected based on low 50% effective inhibitory concentration (EC_50_) values and were produced and purified in-house (**S4 Fig**). None of these candidates inhibited infection with pseudotyped, single cycle VSV particles displaying the SARS-1 S protein (VSV-SARS-1-GFP), indicating specificity for SARS-2 S protein (**S4 Fig. E**). Three of the 16 V_H_H-huFc antibodies, SP1B4, SP1D9, and SP3H4, were identified as showing the greatest potency in viral neutralization with EC_50_s of 0.33, 0.45, and 0.14 nM respectively, against VSV-SARS-2-GFP and EC_50_s of 3.14, 1.12, and 0.70 nM respectively, against *WT* SARS-2 by a plaque neutralization assay (**Fig. 2E-F**). The neutralization values were obtained with three independent virus-based assays (**S5 Table)**, suggesting that SP1B4, SP1D9, and SP3H4 are potent neutralizers of SARS-2, and this observed potency correlates with their strong competition with ACE2 for binding to SARS-2 S protein (**Fig. 2D**). The thermal stability of V_H_H-huFc antibodies was evaluated by differential scanning fluorimetry (DSF). Melting temperatures for SP1B4, SP1D9, and SP3H4 were 60.4°C, 64.0°C, and 61.0°C respectively and are comparable to previously reported SARS-2 neutralizing V_H_H-huFc antibodies (**Fig. 2J; S5 Fig**.).(25)

Dissociation constants were determined for the V_H_H-huFc antibodies using biolayer interferometry (BLI) revealing high affinity for the SARS-2 RBD of 39.5 nM and 8.9 nM for SP1B4 and SP1D9 respectively (**Fig. 2G-H**). SP3H4 exhibited biphasic association and dissociation kinetics which could not be reconciled with a 1:1 global fit analysis and it was therefore not given kinetic parameters (**Fig. 2I)**.

### Epitope Mapping by Escape Mutant Formation and BLI

Epitopes were mapped by sequencing VSV-SARS-2-GFP virus containing escape mutants generated in the presence of SP1B4, SP1D9, or SP3H4, followed by confirmation with BLI binding studies of escape mutant SARS-2 RBDs (**Fig. 3**). Within 48-60 hours of the first passage, viral escape was apparent from all V_H_H-huFc antibodies at the highest concentration tested, with all cells expressing GFP. The viral supernatant used for the second passage was completely resistant to neutralization from a 15-fold excess of the corresponding V_H_H-huFc (**Fig. 3C**). Escape mutations were characterized by RNAseq for each of the VSV-SARS-2-GFP supernatants (**Fig. 3D**). Interestingly, for SP1B4 and SP1D9, E484K (the mutation found in the variant B.1.351 first isolated in South Africa) seems to be selected against in the second passage and Q493R and S494P seem to stabilize as the predominant mutations observed (**Fig. 3D**). This may indicate these two V_H_H-huFc antibodies have similar binding modes. For SP3H4, the L452R mutation seems to be the predominant mutant observed, with none of the virus showing any of the same mutations observed for SP1B4 and SP1D9. This may indicate that this V_H_H-huFc engages the RBD region differently. The L452R mutation is the predominant mutation found in the recently described variants CAL.20A, derived from clade 20A (lineage B.1.232,) and CAL.20C, derived from clade 20C (lineage B.1.429), that are associated with multiple outbreaks in California.(26-28)

**Fig. 3:**
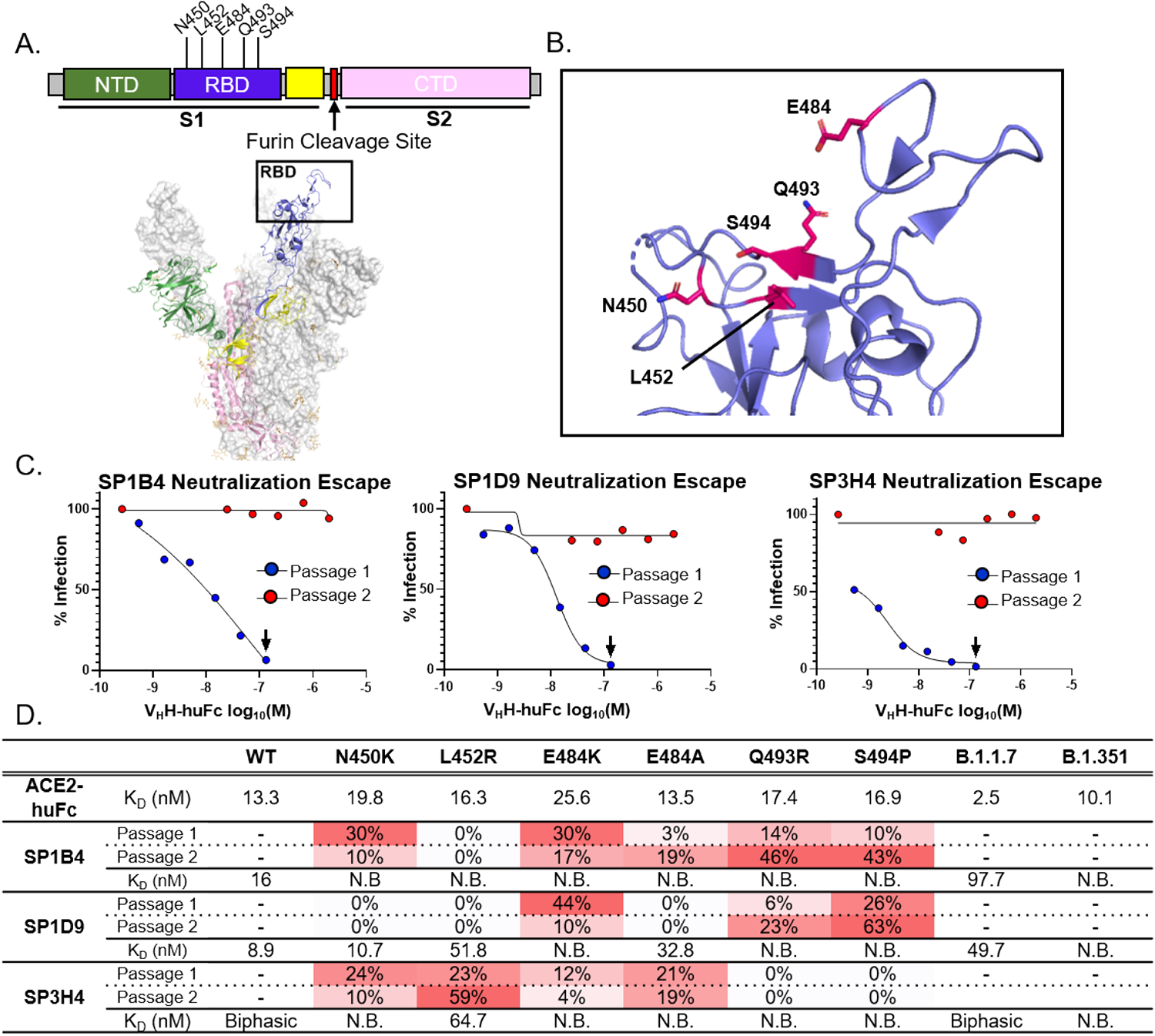
Genereation and Characterization of VHH Neutralization Escape Mutants. **A)** Subdomains of the SARS-2 S gene are color coded with correspoding structural context (PDB: 7CAK). Loci of mutations on S gene from NGS evaluation of virus supernatant from virus infected cells passaged in the presence of V_H_H-antibodies. In this structure the RBD is in the “up” position. **B)** The RBD is designated with the dominant mutations characterized in this study (magenta). **C)** VSV-SARS-2-GFP neutralization escape occurs in a single passage. VSV-SARS-2-GFP neutralization is recorded 12 hpi. The black arrow designated the well from which the virus-containing supernatant was taken and diluted for the second passage. Data for this is from a single replicate. **D)** NGS and kinetic summary of each of the neutralization escape mutant profiles for SP1B4, SP1D9, and SP3H4. Dissociation constants for each individual RBD mutation with ACE2-huFc and V_H_H-huFc antibodies. N.B. = no binding observed at 150 nM SARS-2 RBD by BLI (**S7-9 Fig.**). Sensograms were designated biphasic if they did not agree with a global 1:1 fit analysis. The prevalence of each escape mutation is presented as a percentage at that amino acid postion and is conditionally colored with increasing intesity of red.

SARS-2 RBD variants with single mutations were produced and their affinity for human ACE2 and the three V_H_H-huFc antibodies was determined by BLI (**S6-9 Fig. and S6 Table**). In all cases, individual mutations were shown to have little to no impact on ACE2 binding, suggesting minimal impact on viral infection. Mutations in the RBD significantly impacted V_H_H-huFc binding. Interestingly, SP3H4 maintained the ability to bind to L452R despite this being the predominant mutation observed in the escape mutant VSV-SARS-2-GFP virus generated. Additionally, BLI results show 1:1 binding kinetics for SP3H4 binding to the L452R mutant RBD. The biphasic kinetics observed with SP3H4 and *WT* RBD combined with the 1:1 binding kinetics of SP3H4 for RBD with the L452R mutation may indicate that SP3H4 has multiple binding sites. SP1D9 maintained the ability to bind many of the point mutations generated, though binding was abrogated by Q493R and S494P (**Fig. 3D**) as expected based on the escape mutant analysis. All mutations generated in RBD abrogated binding by SP1B4.

Next, SARS-2 RBDs from recently identified and circulating SARS-2 variants (B.1.1.7 first isolated in the UK and B.1.351) were characterized for affinity to the human ACE2 receptor and top V_H_H-huFc antibodies identified in this study (**S6-9 Fig. and S6 Table**). The B.1.1.7 variant RBD showed an ∼5-fold increased affinity to the ACE2 receptor (K_D_ = 2.5 nM vs 13.3 nM for *WT*). While SP1B4 (K_D_ = 97.7 nM) and SP1D9 (K_D_ = 49.7 nM) maintained binding affinity for the B.1.1.7 RBD, there was a 2.5 -fold and 5.5 -fold decrease in affinity in comparison to the Wuhan RBD (K_D_ = 39.5 nM, and 8.9 nM respectively, (**S7-8 Fig**.) and a visible difference in binding to this RBD for SP3H4 (**S9 Fig**.). For the SARS-2 RBD from the B.1.351 variant, the K_D_ for ACE2 was similar to the K_D_ for the Wuhan strain but there was no detectable binding for SP1B4, SP1D9 or SP3H4 (**S7-9 Fig**.).

### Preclinical Evaluation of V_H_H-huFc antibodies

Top performing V_H_H-huFc antibodies SP1B4, SP1D9 and SP3H4, were assessed for *in vivo* prophylactic (−24 hours) and therapeutic (+24 hours) efficacy against fully virulent *WT* SARS-2 using the recently described K18-hACE2 transgenic mouse model of SARS-2 infection.(29) Two of the three V_H_H-huFc antibodies tested, SP1D9 and SP3H4, provided very significant protection (p < 0.0001) over the isotype control in prophylactically dosed mice. Although SP1B4 was a potent neutralizer *in vitro* it did not demonstrate efficacy against challenge with SARS-2 *in vivo* (**Fig. 4A and 4B)** and was not pursued in further *in vivo* studies. This result highlights the importance of animal validation studies to triage top candidates for clinical evaluation. The SP1D9 group showed 94% survival (n=16), and the SP3H4 group showed 87.5% survival (n=16) at ten days post infection, indicating effective neutralizing activity *in vivo (***Fig. 4A)**. While 81.3% (n=16) of control treated mice lost more than 20% of their starting weight by 6 days post infection (dpi), requiring euthanasia, those pretreated with SP1D9 lost an average of only 3.9% (+/-4.8%) of their starting weight (**Fig. 4B and S10 Fig**.). Similarly, the SP3H4 dosed group exhibited a 5.4% average (+/-9.3%) weight loss. Animals dosed prophylactically with the poorest performing V_H_H-huFc, SP1B4, lost an average of 17.7% (+/-8.3%) of their starting weight, with 50% of animals succumbing to infection by 6 dpi (**Fig. 4B & S10 Fig**.).

**Fig. 4:**
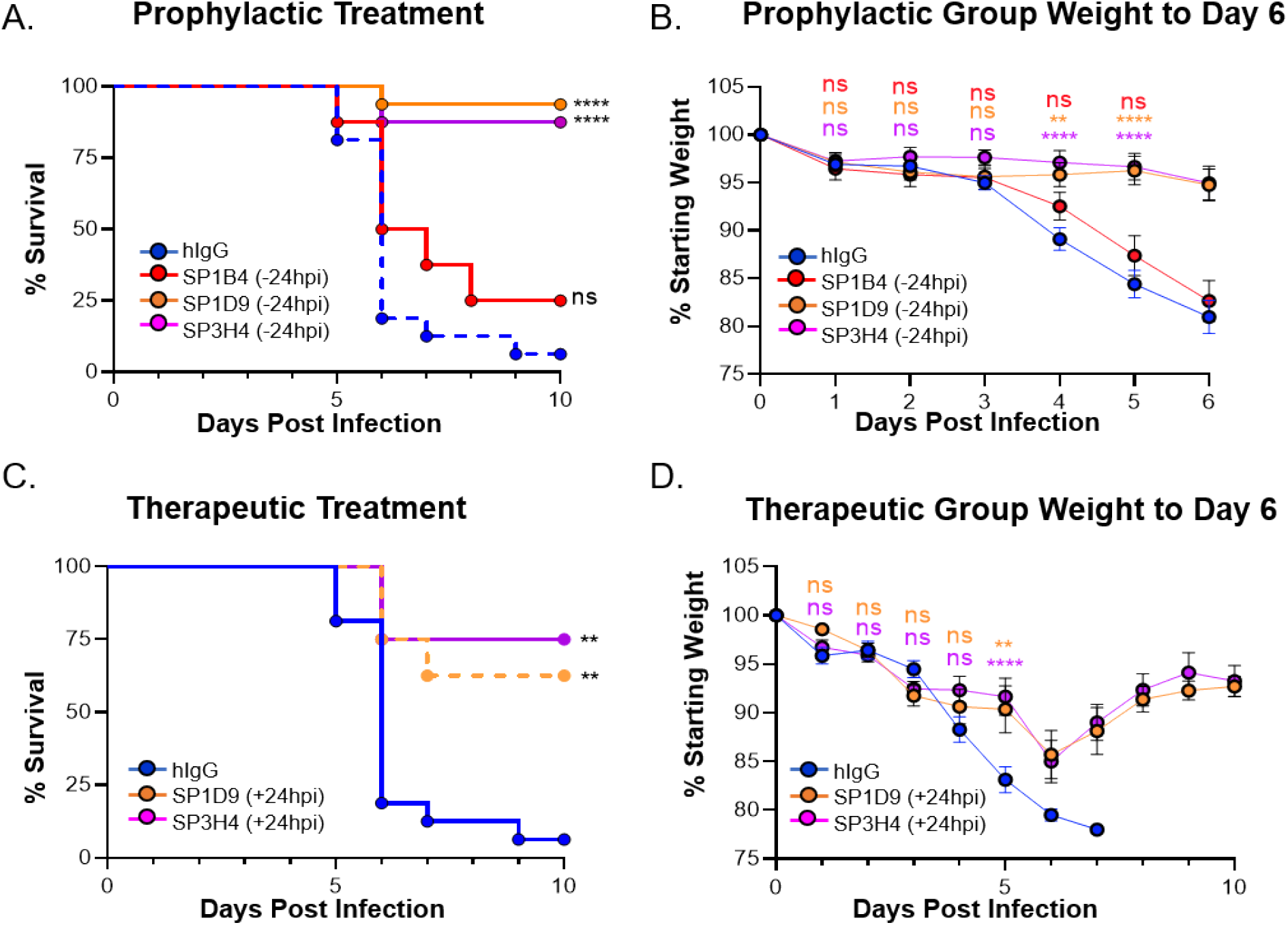
Top V_H_H candidates provide protection from lethal SARS-2 infection *in vivo*. Kaplan-Meier curve illustrating percentage survival of K18-hACE2 mice infected intranasally with 2.5 × 10^4^ PFU SARS-2 and dosed with 10 mg/kg V_H_H-huFc via intraperitoneal injection prophylactically **(A)** at 24 hours prior to infection or therapeutically **(C)** at 24 hpi. A log rank test was performed with Bonferroni multiple comparison correction applied. B) Percentage weight loss for mice in (A). D) Percentage weight loss for mice in (C). For **(B)** and **(D)**, statistical analyses were performed at time points when all mice were alive to avoid survivor bias. Data displayed are the mean +/- standard error of the mean. **A** & **B**) Isotype control n=16, SP1B4 (−24 hpi) n=8, SP1D9 (−24 hpi) n=16, SP3H4 (−24 hpi) n=16. **C** & **D**) Isotype control n=8, SP1D9 (+24 hpi) n=8, SP3H4 (+24 hpi) n=8. Data represent two experiments (**** p < 0.0001; ** p < 0.01; ns=not significant).

Since SARS-2 viral titers peak as early as 48 hours post-infection (hpi) *in vivo*, allowing the virus to establish infection for 24 hours prior to therapeutic administration should demonstrate the versatility of these V_H_H-huFc antibodies in disease treatment.(29) To evaluate the efficacy of SP1D9 and SP3H4 V_H_H-huFc antibodies post-exposure, mice were infected intranasally with 2.5×10^4^ PFU SARS-2, then administered 10 mg/kg of V_H_H-huFc antibodies 24 hpi. Impressively, both SP1D9 and SP3H4 demonstrated significant therapeutic protection (p = 0.0033, p = 0.0013, respectively) over that of the isotype control. The SP1D9 group showed 58% survival (n=8), and the SP3H4 group showed 75% survival (n=8) at ten dpi (**Fig. 4C)**. Of note, several surviving individuals in each of the post-exposure V_H_H-huFc-treated groups exhibited 10-15% weight loss up to 5 dpi, but then rebounded from 6-10 dpi, gaining back 90-95% of their starting weight (**Fig. 4D** and **S10 Fig**.). Taken together, these data demonstrate protection from lethal infection after a single 10 mg/kg dose, and further suggest that neutralization of SARS-2 by V_H_H-huFc antibodies *in vivo* can promote recovery from an ongoing infection.

## Discussion

The COVID-19 pandemic caused by SARS-2 has resulted in 146 million cases with 3.09 million deaths and an estimated global economic cost of over 10.3 trillion dollars in forgone output for 2020 and 2021.(30) As variants of SARS-2 emerge, it is essential to possess a diverse set of prophylactic and therapeutic tools, in order to maintain the progress made toward ending this pandemic. While vaccines have been great successes, therapeutic biologics are emerging as a critical tool in preventing progression to severe disease in those who do become infected.(31)

Several therapeutic antibody candidates with efficacy against SARS-2 have been recently identified, including three with emergency use authorization, but there are a number of caveats associated with conventional antibodies as therapeutics such as time-consuming discovery, relying on immunized or patient sera, expensive and labor-intensive production, and the large doses required to achieve clinical efficacy. (9, 22, 32-34) High stability, solubility, and the ability to be multimerized, are just some of the many reasons why single-domain heavy chain-based antibody therapeutics (V_H_H-Fc) represent a highly promising method for treatment.(12) One V_H_H-based therapeutic is already approved by the FDA for treatment of a rare blood clotting disorder and many more are in late stage clinical trials.(12, 35) The small size of V_H_H antibodies and the wider distribution of CDR3 loop lengths compared to human IgGs expands the type of epitope which can be effectively targeted.(12) Finally, preliminary evidence suggests that V_H_H-Fc antibodies are transported more efficiently into the blood and lung parenchyma following intraperitoneal administration compared to conventional IgG1 antibodies.(36, 37) Several groups have recently used *in vitro* screening techniques to identify V_H_H domains with high affinity for the SARS-2 S protein and neutralization of SARS-2 with EC_50_s ranging from 0.02 nM to 2 μM in pseudotyped virus and in *WT* SARS-2 virus assays.(35, 38-45) However, in addition to neutralization, there are several antibody and immune functions that contribute to prophylactic and therapeutic effectiveness *in vivo*, and only one of these studies characterized the *in vivo* efficacy of a single V_H_H-Fc construct (V_H_H-Fc ab8) at reducing lung viral titers at 2 or 5 days post infection.(36) Importantly, the mouse-adapted SARS-2 and the golden hamster SARS-2 infection models used in this study are not lethal, rather, they inform efficacy for more mild disease cases.(29, 36, 46, 47) In this regard, the K18-hACE2 SARS-2 infection model is particularly valuable for testing the protective and therapeutic properties of novel antibodies in the context of severe infection and disease.(48, 49)

In this study, we have identified and characterized several prospective therapeutic neutralizing nanobodies (V_H_Hs) both *in vitro* and *in vivo* from a high diversity synthetic library. Three candidate V_H_H-huFc antibodies are highlighted in this study because of their potent sub-nanomolar EC_50_s in preventing viral infection. For the SP1B4, SP1D9, and SP3H4 huFc constructs, affinity maturation was not required to achieve potent neutralization against SARS-2 pseudovirus (EC_50_s of 0.33, 0.45, and 0.14 nM, respectively) or *WT* SARS-2 virus (EC_50_s of 3.14, 1.12 and 0.70 nM, respectively). Two of the V_H_H-huFc antibodies provided protection of K18-hACE2 transgenic mice when administered 24 hours before infection with *WT* SARS-2. A single dose of SP1D9 or SP3H4 exhibited significant therapeutic value when administered a full 24 hpi, which is closer to the peak of infection than was used to evaluate therapeutic efficacy of V_H_H-Fc ab8 (6hpi).(36, 48) In the present study, the onset of weight loss at 2-4 dpi, indicative of effective disease progression, and subsequent improvement after treatment establishes the potential for efficacy in the presence of an established SARS-2 infection (**Fig. 4D and S10 Fig**.).

Selection for escape mutants by individual V_H_H-huFc antibodies demonstrate the high propensity for SARS-2 to escape neutralization *in vitro*. Several studies, including this one, have generated mutations *in vitro* that exist in circulating SARS-2 variants and revealed that these mutations partially or completely block binding of top anti-SARS-2 therapeutic antibodies.(50) The potential for continued mutation acquisition and escape in SARS-2 highlights the critical importance of having multiple options for antibody cocktails.(39) Furthermore, sequencing analysis of the latest strains’ mutation status would allow patient stratification to identify which therapeutic regimen will be most effective in combating the latest emerging variant.(39)

In this study, viral escape was readily apparent in the presence of our top candidates, with several mutations in the RBD. Interestingly, many of the initial mutations were selected against in the second passage (**Fig. 3**) and none of these individual mutations in the RBD significantly impacted binding to ACE2 (**Fig. 3D and S6 Fig**.). This suggests that these positions of the S gene are highly susceptible to single-nucleotide polymorphisms and should be considered when identifying future preclinical antibody candidates as they may be present in future variant strains. Escape mutations observed for our two most promising candidates are all found in proximity to each other likely indicating overlapping epitopes, though they may engage this epitope with different geometries (**Fig. 3**). All three V_H_H antibodies neutralize viral infection by targeting the RBD of the SARS-2 S protein and compete directly with ACE2 binding to trimeric SARS-2 S which indicates they are class I nanobodies.(35) Although the V_H_H antibodies described in this study bind to a single epitope, other groups have recently identified V_H_H antibodies that target a distinct epitope and could be used in combination with the V_H_H antibodies identified in this study to develop highly potent therapeutic cocktails that prevent the emergence of escape mutants.(39, 41, 45)

We have also evaluated our V_H_H-huFc antibodies’ ability to engage clinically relevant SARS-2 variants (B.1.1.7 and B.1.351) which pose a significant hurdle to therapeutic efficacy and vaccine-derived immunity. Interestingly, we observed a ∼5-fold increase in affinity of the B.1.1.7 variant RBD for the ACE2 receptor, which may account for the increased infectivity observed with this strain of SARS-2. While none of our most promising candidates bind to the B.1.351 variant, SP1D9 and SP3H4 maintained binding affinity to the B.1.1.7 RBD variant (**S8-9 Fig**.). The E484K mutation, observed in the B.1.351 variant and now a new variant B.1.526, confers total abrogation of V_H_H-huFc binding. Another clinically relevant mutation, L452R, which can be found in circulating variant strains CAL.20A and CAL.20C, was selected for with SP3H4. Interestingly, SP3H4 retains affinity for SARS-2 RBD with the L452R mutations. Furthermore, the kinetics appear to shift from biphasic to 1:1 with the L452R mutation, which may indicate that SP3H4 has two non-overlapping binding sites. While many of the mutations in clinically relevant SARS-2 variants emerged in this study, there are several mutations that arose in this study which have yet to be highlighted. Using these antibodies to rapidly generate escape mutants could help to preemptively identify and develop countermeasures for future variants before they arise and circulate in the population.

In this work we have reported the design of a large synthetic V_H_H library (3.18 × 10^10^) and described the library’s utility in identifying over 50 V_H_H candidates which bind to SARS-2. We triaged top candidates based on neutralization efficiency *in vitro*, and evaluated efficacy *in vivo* using a mouse model of severe SARS-2 disease. To our knowledge, this is the first study to examine survival up to ten dpi with a single dose of V_H_H-Fc given 24 hpi in the context of a lethal SARS-2 infection. While *in vitro* neutralization assays and short-range *in vivo* titer reduction studies are informative, they do not always recapitulate likelihood of survival. There are several reasons why *in vitro* potency may not correlate with *in vivo* protection, including an antibody’s pharmacokinetic properties and/or a broad range of immunomodulatory functions required for viral clearance.(49, 51-53). Studies are currently underway to evaluate the efficacy of V_H_H-huFc combinations administered at later time points post-infection and to investigate the role of Fc effector function in protection against SARS-2. The demonstrated ability of this virus to generate escape mutants emphasizes the need for multiple therapeutic options, with demonstrated *in vivo* activity in a post-infection context, to rapidly react to the emergence of resistant variants as expanded vaccine and antibody therapy use creates selection pressures on the viral population.

## Acknowledgements

We would like to thank Prof. Sean Whelan from Washington University School of Medicine St. Louis for graciously providing access to the pseudotyped VSV-SARS-GFP virus used in this study. We would also like to thank Dr. Pei Yong Shi from the World Reference Center for Emerging Viruses and Arboviruses at University of Texas, Medical Branch for generously providing the infectious clone of SARS-CoV-2 expressing a NeonGreen reporter gene. We would also like to thank Robert Meager for critically reviewing the paper. This work was supported by the Laboratory Directed Research and Development Program at Sandia National Laboratories and the Department of Energy (DOE) Office of Science through the National Virtual Biotechnology Laboratory, a consortium of DOE national laboratories focused on response to COVID-19, with funding provided by the Coronavirus CARES Act. Sandia National Laboratories is a multi-mission laboratory managed and operated by National Technology & Engineering Solutions of Sandia, LLC, a wholly owned subsidiary of Honeywell International Inc., for the U.S. Department of Energy’s National Nuclear Security Administration under contract DE-NA0003525. This paper describes objective technical results and analysis. Any subjective views or opinions that might be expressed in the paper do not necessarily represent the views of the U.S. Department of Energy or the United States Government. All work performed at Lawrence Livermore National Laboratory is performed under the auspices of the U.S. Department of Energy under Contract DE-AC52-07NA27344.

## Supporting Information Captions

**S1 Fig**. Distribution of the lengths of CDR loops 1-3 of 670 VHH sequences deposited in the Single Domain Antibody Database (sdAB-DB).

**S2 Fig**. Monoclonal Phage-VHH screening for SARS-2 S and RBD specific binding clones: ELISA with 384 clonal VHH-displaying phage preparations with SARS-2 S (left), SARS-2 RBD (center), and BSA (right).

**S3 Fig**. Triage of VHH-huFc candidates: A) ELISA with 54 VHH-huFc with SARS-2 S (left), SARS-2 RBD (center), and BSA (right). The data are from experimental conditions performed in triplicate, the error is the standard deviation from the mean.

**S4 Fig**. Reconfirmation of Purified VHH-huFc candidates: A) ELISA with 16 VHH-huFc antibodies with SARS-2 S. B) Binding of 16 VHH-huFc candidates to SARS-2 RBD. C) Competition ELISA with the VHH-huFc antibodies. D) Neutralization of VSV-SARS-2-GFP infection of Vero cells. MM57 is a control anti-SARS-2 neutralizing monoclonal antibody from Sino Biological. The anti-VSV monoclonal antibody was produced from hybridoma line CRL-2700 (ATCC). E) VHH-huFc antibodies do not neutralize SARS-1 pseudotyped VSV demonstrating specificity for SARS-2 S. M396 is an anti-SARS-1 antibody produced by GenScript. The data for a-e are from experimental conditions performed in triplicate, the error is the standard deviation from the mean.

**S5 Fig**. Differential scanning fluorimetry to determine VHH-huFc melting temperatures.

**S6 Fig**. BLI Sensograms for ACE2-huFc binding to SARS-2 RBD variants and escape mutants.

**S7 Fig**. BLI Sensograms SP1B4 binding to SARS-2 RBD variants and escape mutants.

**S8 Fig**. BLI Sensograms SP1D9 binding to SARS-2 RBD variants and escape mutants.

**S9 Fig**. BLI Sensograms SP3H4 binding to SARS-2 RBD variants and escape mutants.

**S10 Fig**. Weight change for individual mice from preclinical evaluation of VHH-huFc antibodies.

**S1 Table:** Next-Generation Sequencing Results for VHH Library Diversity

**S2 Table:** Next-Generation Sequencing Results for VHH Library CDR3 Length Distribution

**S3 Table:** Quantification of Phage from Biopanning Campaign against SARS-2 S and RBD

**S4 Table:** Summary of ELISA, Competition ELISA and neutralization of VSV-SARS-2-GFP infection of Vero cells for VHH-huFc’s (Y =Yes, N = No, NT = not tested

**S5 Table:** Summary of Neutralization of VSV-SARS-2-GFP, SARS-2-NG, and SARS-2 infection of Vero cells for reconfirmation VHH-huFc antibodies.

**S6 Table:** Summary of BLI data for escape mutant and SARS-2 variant’s

